# Phenotero: annotate as you write

**DOI:** 10.1101/324053

**Authors:** Daniela Hombach, Jana Marie Schwarz, Ellen Knierim, Markus Schuelke, Dominik Seelow, Sebastian Köhler

## Abstract

Controlled vocabularies and ontologies have become a valuable resource for knowledge representation, data integration, and downstream analyses in the biomedical domain. In precision medicine, especially in clinical genetics, the Human Phenotype Ontology (HPO) as well as disease ontologies like the Orphanet Rare Disease Ontology (ORDO) or Medical Subject Headings (MeSH) are often used for *deep phenotyping* of patients and coding of clinical diagnoses. However, the process of assigning ontology classes (annotating) to patient descriptions is often disconnected from the process of writing patient reports or manuscripts in word processing software such as Microsoft Word or LibreOffice. This additional workload and the requirement to install dedicated software may discourage usage of ontologies for parts of the target audience.

To improve this situation, we present *Phenotero*, a freely available and simple solution to annotate patient phenotypes and diseases at the time of writing clinical reports or manuscripts. We adopt Zotero, a well-established, actively developed citation management software to generate a tool which allows to reference classes from ontologies within clinical reports or manuscripts at the time of writing. We expect this approach to decrease the additional workload to a minimum while ensuring high quality associations with ontology classes. Standardised collection of phenotypic information at the time of describing the patient allows for streamlining of clinic workflow, efficient data entry, and will subsequently promote clinical and molecular diagnosis, remove ambiguousness from manuscripts, and allow sharing of anonymised patient phenotype data with ultimate goal of a better understanding of the disease. Thus, we hope that our integrated approach will further promote the usage of ontologies and controlled vocabularies in the clinical setting and in the biomedical domain.

## Introduction

Information driven approaches are becoming ever more important in biological and medical science. Consequently, biomedical ontologies are used in several research projects and applications ^1–4^. Ontologies are often defined as a formal specification of a shared conceptualisation^5^. In research and in the clinic, these ontologies provide a standardised, machine-interpretable way to describe complex biomedical concepts such as human clinical features, animal phenotypes, or gene properties. The application of ontologies is essential for robust information retrieval but also for conducting downstream analyses in the study of diseases, e.g. for differential diagnostics and exploration of pathomechanisms^6–8^.

The Human Phenotype Ontology (HPO^9,10^), for instance, is widely used to collect and characterise the symptoms and clinical features of a patient by referencing standardised ontology terms (HPO classes) for individual phenotypic features. To encode diagnoses, other ontologies like Medical Subject Headings (MeSH, www.nlm.nih.gov/mesh/), Disease Ontology (DO^11^), or the Monarch Disease Ontology (MONDO, www.obofoundry.org/ontology/mondo.html) are often used.

While the HPO has become the standard to describe patients’; phenotypes in genetic databases and to exchange phenotypic data, medical reports still suffer from a lack of standardisation. The journal *Cold Spring Harbor Molecular Case Studies* has hence already started to require HPO-based descriptions in manuscripts, thus making patients’; descriptions easily recordable, exchangeable, and comparable with other publications, studies, and database contents.

The association between an ontology class (e.g. *short stature*) to an object (e.g. a patient) is often termed *annotation* and requires expert curation. In medicine, physicians have to translate their plain medical terms into standardised ontology entries. This is a highly specialised task, which rises and falls with the physician’s expertise both regarding clinical features and the use of ontologies. Current methods used for medical text are dedicated annotation software solutions such as *PhenoTips*^12^ and Patient Archive (patientarchive.org). Another approach is to retrospectively recognise ontology classes in written text using text mining software, for instance NCBO annotator^13^ or BioLarK^14^.

*PhenoTips* can be used to generate a local patient database and to annotate patients with terms from the HPO. However, such software adds another layer of complexity as they are disconnected from the writing process and require to step away from word processing tools in order to identify the best matching ontology classes. In addition, they often require non-trivial software installation: *PhenoTips*, for instance, is optimised to run on a dedicated Linux server, creating a nearly insurmountable obstacle in many clinical work environments. The test version of *PhenoTips* is available for Windows computers, but is not recommended for production use. Moreover, these tools are aimed at much more complex tasks like the representation of growth charts or pedigree information and are not ideally suited for the mere assignment of ontology-based terms to human phenotypes in a manuscript or clinical report.

Text mining software, on the other hand, is usually run retrospectively and hence also disconnected from the writing process. Moreover, NCBO Annotator or BioLarK require the text to be sent via the internet to an external server, which can cause data security issues, especially when dealing with confidential patient information. This poses a severe problem when it comes to medical reports which routinely include personal and identifying information as names, addresses, and diagnoses. Further, automatic solutions often try to assign ontology classes to all terms or phrases that vaguely resemble a human phenotype, leading to faulty and confusing annotations. Moreover, these tools frequently fail in distinguishing actual patient-related phenotypes from terms related to family history. Tools such as NCBO annotator are also problematic when it comes to negated mentioning of concepts. This makes the resulting data noisy and usually requires subsequent manual curation.

Our goal is to close the gap between writing and annotating to enable physicians and clinical and biomedical researchers to seamlessly associate standardised biological or clinical terms as they write: Our free tool *Phenotero* allows referencing ontology classes from HPO and MONDO directly within word processing software such as Microsoft Word or LibreOffice (*annotate-as-you-write*). Hence, authors of clinical reports or manuscripts can link patient symptoms or diseases to an ontological entry, e.g. from the HPO, while they are composing their text; much like adding a reference to a scientific paper. This tremendously eases later usage of information buried in the text because it safes downstream users to conduct the association from symptoms to ontology classes over and over again. Moreover, it enables the person who knows the patient best-the physician or researcher familiar with their case-to annotate phenotype terms without delay, leading to high-quality associations.

*Phenotero* is based on the free, open-source citation software Zotero and thus builds on existing features: a word-processor plugin, standardised and extendible citation styles, a search function, and active community support.

## Methods

### Conversion of HPO and MONDO to libraries

Our primary goal was to keep the user within the Zotero application without the necessity for further external tools. In order to import the HPO and MONDO ontologies into Zotero we provide two strategies:

The **first** and simpler method is to join the phenotypes and diseases group on the Zotero website (see www.zotero.org/groups/2168222/phenotypes. and www.zotero.org/groups/2168493/diseases.). Once the user has joined these groups, they are automatically being notified about updates in the database file, which can then be imported to their Zotero working environment with just one click (Zotero sync function).

A **second** option is the import via JSON files which we make available for download (https:phenotero.github.io/data_json/.). For this, we transferred the OBO versions of the ontologies into JSON-format (HPO: releases/2018-03-08, MONDO: releases/2018-03-16). Within Zotero, users can then import these files into the library of their choice. The obo2json converter is written in Java and can be obtained from GitHub (https://github.com/phenotero/obo2json.). In a digital bibliography, entries must have values for specific fields such as title, author and type. For *Phenotero*, each ontology class is defined as an “entry-dictionary” type, to better distinguish it from articles, books, etc. This reference-type is rarely used in biomedical articles and should hence not interfere with other Zotero capabilities. The ontology IDs are stored in the container-title field and the synonyms under author, as this field allows to have multiple entries per item. The exact mapping between OBO fields and JSON-format for Zotero is described in Table 1. Note that we converted the ontologies to JSON-format in order to create the group libraries described before.

We decided to use MONDO, as this ontology integrates a major fraction of the most important disease ontologies such as MeSH, DO, ICD9, ICD10, NCI Thesaurus, OMIM, and Orphanet.

**Table 1:**
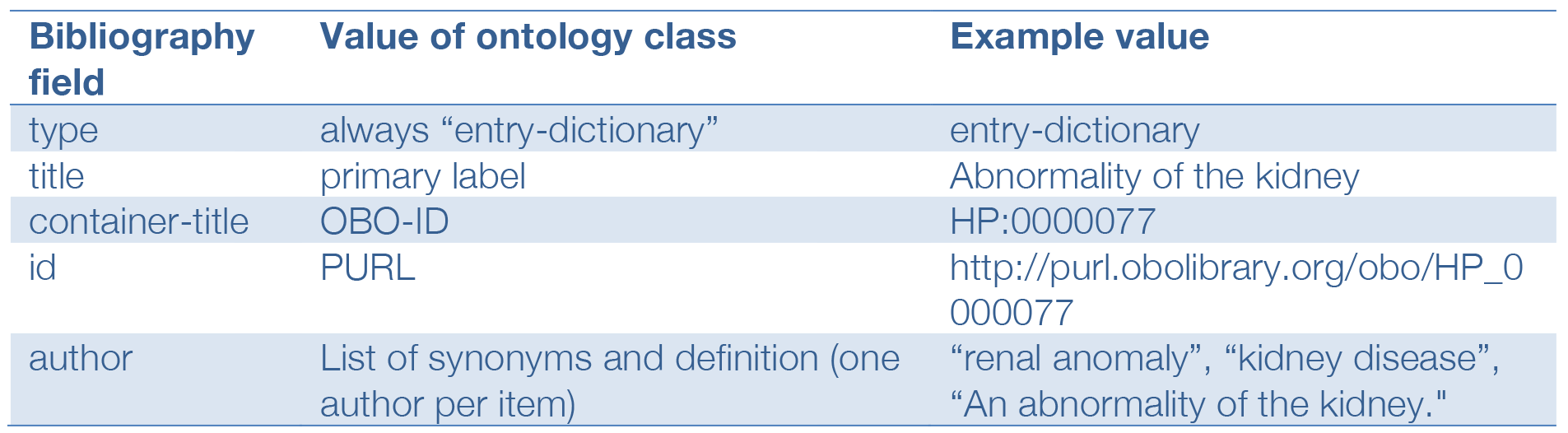
Mapping description for creating the JSON-format version of an OBO ontology class. The resulting JSON-file was used for the initial import of the ontologies into Zotero. Code for the OBO-to-JSON conversion is available at and https://github.com/phenotero/obo2json

| Bibliography field | Value of ontology class | Example value |
| --- | --- | --- |
| type | always "entry-dictionary" | entry-dictionary |
| title | primary label | Abnormality of the kidney |
| container-title | OBO-ID | HP:0000077 |
| id | PURL | http://purl.obolibrary.org/obo/HP_0000077 |
| author | List of synonyms and definition (one author per item) | "renal anomaly", "kidney disease",<br>"An abnormality of the kidney." |

### Definition of Phenotero citation styles

The style of the actual citations and bibliographies or reference lists in the final document is determined by style sheets written in the citation style language (CSL). For most scientific journals a citation style in CSL format is available, which can be easily imported into Zotero and afterwards selected within the word processing plugin.

To provide convenient ways of incorporating *Phenotero* citations and reference lists into word processing software such as Microsoft Office Word, we defined various citation styles: Two stand-alone files, “Phenotero-Phenotypes” (phenotype.csl) and “Phenotero-Phenotypes-IDlist” (phenotype_idlist.csl) are applicable for users who only wish to add phenotype annotations without generating a common bibliography. The style “Phenotero-Phenotypes” lists the Ontology class IDs together with their primary label, e.g., “[HP:0000767] Pectus excavatum”. “Phenotero-Phenotypes-IDlist”, which is intended for further downstream use in external tools, only shows the IDs themselves, so that they can be easily copied into other tools.

In addition, we provide a number of style sheets which allow incorporation of phenotype annotations into commonly used journal citation styles. These styles put all articles and other library objects into a standard reference list, but append the references of phenotype and disease ontology classes at the end. In the future, further journal citation styles could be adapted in a similar fashion. All *Phenotero csl*-files are available for download at phenotero.github.io/data_csl.

## Results

*Phenotero* is a simple tool for referencing phenotypes and diseases in written text and allows physicians and clinical researchers to readily stay within word processing software. This enables a novel workflow for direct in-text annotations of ontology classes, which we call “*annotate as you write*”.

### Setting up Phenotero

*Phenotero* requires a running installation of Zotero and the associated Microsoft Office Word or LibreOffice Writer plugin. These plugins are bundled with Zotero from Version 4.0 upwards and add a Zotero toolbar to Word or Writer. Zotero runs on Windows, Mac, and Linux operating systems and the installation is easy and quick (www.zotero.org.).

Following this, users should join the *Phenotero* Zotero Groups (www.zotero.org/groups/2168222/phenotypes. and www.zotero.org/groups/2168493/diseases.). Here, they can obtain the latest MONDO and HPO libraries. Zotero will automatically start to synchronise contents of the groups to the local machine. The current *Phenotero* MONDO library contains 20,503 diseases and it takes around 40 minutes to import it into Zotero. We omitted MONDO text definitions from this library in order to reduce time required for import. The current HPO library contains 13,342 phenotypes and is imported within 20 minutes.

The citation style files in CSL format are available at phenotero.github.io/data_csl and can be imported within seconds. We currently offer seven citation styles, but will make more available upon user request. More information about *Phenotero’s* installation and usage can be found in the Supplementary Material as well as online on the website at phenotero.github.io.

### Use Cases

Three simple case examples shall help to highlight the benefit of *Phenotero’s*usage.

In the **first scenario,** the phenotype information from a clinical report or manuscript can be extracted for data exchange or further analysis with an HPO-based tool, such as PhenGen^15^, eXtasy^16^, PhenIX^2^, or MutationDistiller. The latter is especially handy in the case of software that lacks auto completion to suggest suitable ontology classes matching the entered text. In this scenario, users need to select the *Phenotero* citation style “Phenotero-Phenotypes-IDlist” and copy-paste the list of HPO-IDs from the bottom of the text into one of the aforementioned tools (see Figure 2 centre).

In the **second scenario,** a physician or researcher is writing a report about a counselling to a patient. Beyond describing their disease and clinical symptoms, the corresponding sentences are then annotated with HPO references (see Figure 2 left). A recent study has presented that patients appreciate very much a proper description of their medical condition at eye level^17^. Having links to ontologies such as HPO or MONDO helps the patient to find textual definitions, lay person descriptions, and further information about the mentioned phenotypes and diseases.

**Figure 1:**
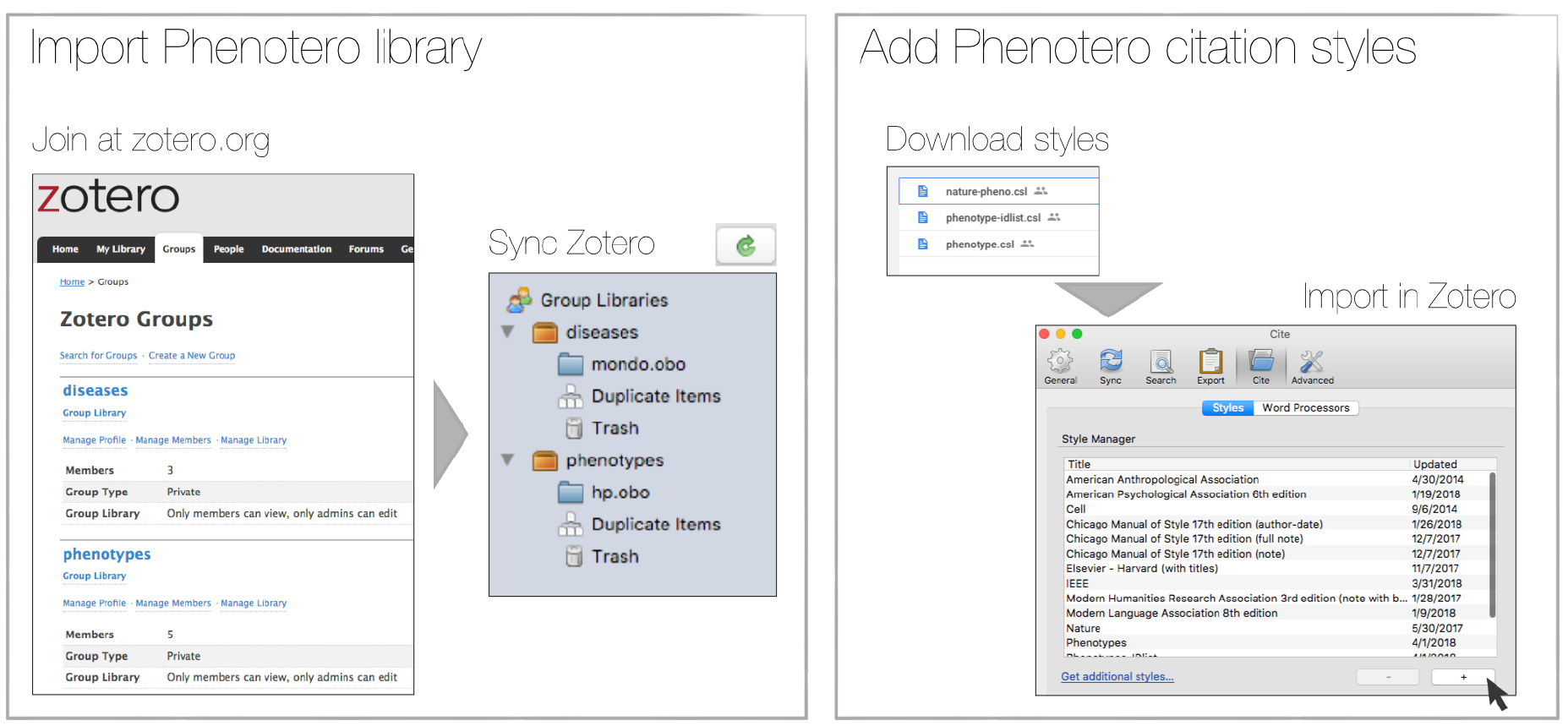
Following the installation of Zotero, users have to perform two steps to perform in order to enable *Phenotero*: Import the *Phenotero* library and add the appropriate citation style which can be retrieved from phenotero.github.io/data_csl.

**Figure 2:**
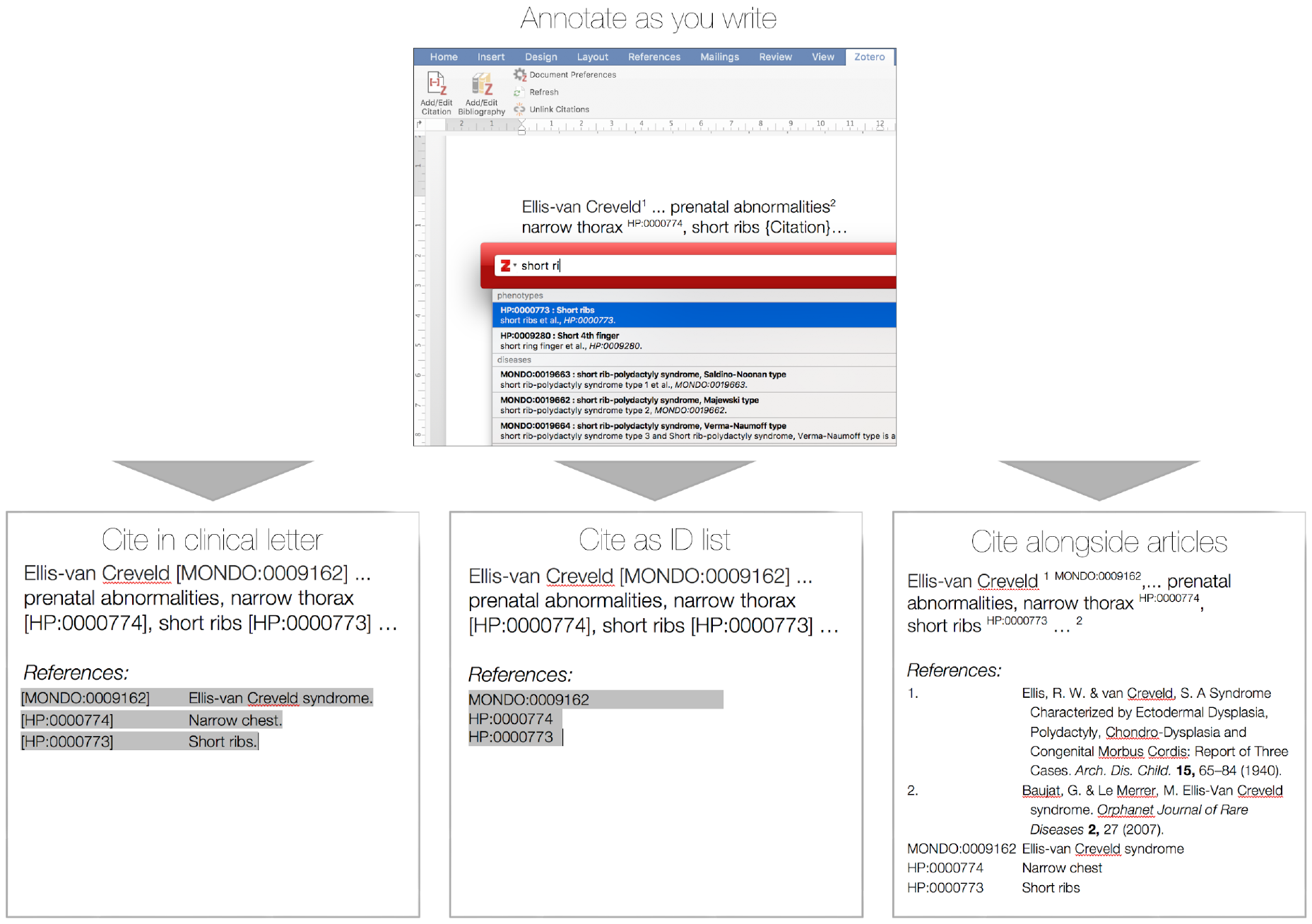
Workflow of *Phenotero’s* “annotate as you write”. Zotero comes with Word and Writer plugins, allowing for the addition of references during the writing process. Once the text has been annotated, different bibliographies can be added.

The **third example** is based on the assumption that an increasing number of journals are seeking ways to publish manuscripts with references to controlled vocabularies. We assume that for this purpose, HPO or MONDO references shall be integrated into the reference section. This is a prototype implementation, but we expect this to be useful for several biomedical journals, especially those that already request HPO encoded phenotypes to be submitted such as *Cold Spring Harbor Molecular Case Studies* or *Orphanet Journal of Rare Diseases*. We provide citation styles for multiple journals (e.g. CSH-Laboratory-Press, AJHG, Orphanet Journal of Rare Disease, and Nature), which will append the referenced ontology classes to the reference section (see Figure 2 right).

In all scenarios, the *Phenotero* approach makes the phenotype and disease information amenable to automated computational analyses of written reports, as the referenced ontology classes can be easily extracted from the text documents. Thus, the framework can also be used to gradually build a patient database consisting of text documents. Text documents become easily indexable, searchable and amenable to sophisticated clustering and similarity calculation algorithms. For this purpose, we created an executable jar file available (see *Phenotero* website at phenotero.github.io) that processes all *docx-* and *odt-*files in a directory and its subdirectories and generates a *tsv*-file with all extracted ontology identifiers per document. This file can then be used for further analyses.

## Discussion

*Phenotero* provides a quick and easy solution for referencing ontology-based phenotypes and diseases by direct in-text annotations of classes of ontologies such as the HPO or MONDO. It is an extension of the Zotero citation management open source software, which runs on all common operating systems and is easily integrated into multiple word processors. Users are confronted with a minimal amount of computational complexity and can readily stay within their usual working environment, e.g. Microsoft Office Word or LibreOffice Writer.

The use of *Phenotero* is very simple and almost identical to writing a manuscript using Zotero, users can find thorough und user-friendly documentation online at www.zotero.org. We additionally provide in-depth, Zotero-specific documentation on our website.

Usage of ontology-based phenotypes in data that is shared with others, such as manuscripts and even physicians’; reports, has many advantages. First of all, a fully described patient’s phenotype leaves less room for uncertainties than full-text descriptions often do. In addition, the phenotypic data becomes computer-readable and can easily be processed in further steps, e.g. added to databases or analysed for similarities. Another issue is privacy: The use of ontology-based symptoms allows easy separation of phenotypic from personal information in medical records. This is very important when data is exchanged-complete medical records contain many items to make a patient easily identifiable. A mere list of ontology terms is obviously stripped from all these data and can hence be shared with others without revealing the patient’s identity. This new way of referencing ontologies in written reports does not only provide an easy-to-use tool but a solution to the important issue of data privacy as no sensitive data has to be uploaded to the internet, e.g. to apply text mining.

A potential drawback is that Zotero allows for only one bibliography per document, which may cause problems if users aim for separate bibliographies for publications and ontology references. For this purpose, we found it useful to offer the option of sorted bibliographies which separate references from ontology annotations (see Figure 2 right).

Although *Phenotero* is currently designed for phenotype and disease ontologies, it can in principle work with any ontology that can be transformed into a digital bibliography. Extensions that enable various annotations from the field of anatomy, cell or gene ontology are principally possible. The *Phenotero* approach is almost language independent, i.e. the clinical reports and manuscripts can be written in any language and still contain references to the ontologies. However, searching the referenced ontology classes requires English, but may be simplified once the ontologies have been completely translated into the target language (see e.g. crowdin.com/project/hpo-translation). Finally, we note that the presented approach is not limited to Zotero but can also work with Endnote (alias: Phendnote), Mendeley (alias: Phendeley), since the libraries, can be exported from Zotero and imported to other reference management software as well. However, we chose Zotero for its ease of use and open-source/free use policy.

With *Phenotero*, we hope to get one step closer towards ubiquitous deep phenotyping in a standardized format. We deem this goal beneficial for various downstream tasks on the side of patients as well as of physicians and researchers.

## Supplement

### Import Phenotero library

There are two methods for importing the *Phenotero* library.

1. The simplest way to ensure up-to-date library access is to join our *Phenotero* groups: When you are logged in with your Zotero account on the Zotero website, join the *Phenotero* groups called “phenotypes” and “diseases” (go to www.zotero.org/groups/2168222/phenotypes. and www.zotero.org/groups/2168493/diseases.). The next time you open your Zotero application (and are logged in with you Zotero account), Zotero will start syncing these libraries under “Group Libraries”. This syncing process will take about 20 minutes for the “phenotypes” and about 40 minutes for the “disease” library.
2. You can obtain JSON files for the phenotypes and diseases libraries from https://phenotero.github.io/data_json. After you have downloaded these files, you can select “File”, “Import”, and then select one of the downloaded files. This will add the libraries to your local library.

### Import csl-files

In order to obtain the citation styles, go to https://phenotero.github.io/data_csl/ and download them. Afterwards, you have to open Zotero and go the *Settings* menu. From there, you select “Cite” and get a list of all currently imported citation styles. You can then add the *Phenotero* styles by clicking on the button labelled with a “+” and then selecting the csl-file you downloaded. You have to repeat the last step separately for each of the styles you wish to import.

### Annotate as you write in word processing software

Referencing a phenotype or disease entry from the *Phenotero* library is done by using the “Add/Edit Citation” button provided by the Zotero plugin within your text processing software. In case that this is your first reference in the text, the plugin will ask you which citation style to use. Choose one of the styles that we have currently made available. Afterwards you can simply reference all phenotypes and disease in your text.

If you want to add a bibliography to your text, simply use the “Add/Edit Bibliography” button within the Zotero plugin.

### Instructions to modify a csl-file

This is an example showing how we extended the nature csl-file to enable *Phenotero* support for displaying *Phenotero* entries together with conventional references in a single document. Please note that this is only feasible if you are not using any other items of type ‘entry-dictionary’ in your reference list. This should usually be the case for journal articles and clinical letters. Please be also aware, that other csl-files might be very different and may require different approaches to enable *Phenotero*–style bibliographies.

### Change file name and title

In the info section, file name and title have to be changed for the new file to function in Zotero:

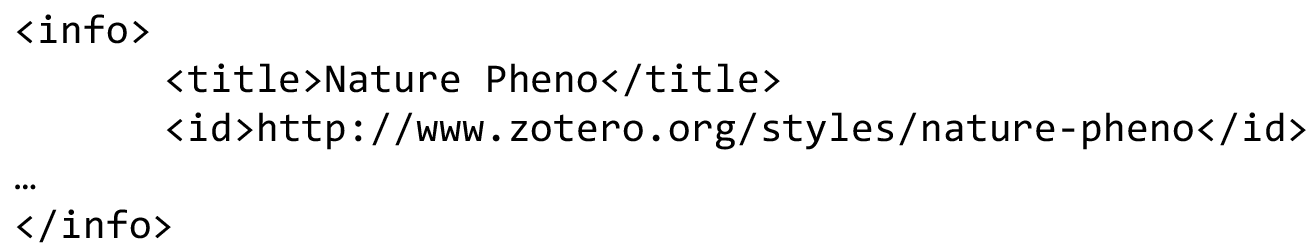

### Add Macros

Next, we added macros to change the default numbering of the citation to the ontology identifiers and to separate the ontology entries from the ‘normal’. These macros can be added anywhere in the MACRO section of the template csl-file. This allows *Phenotero* to display all bibliography references first, followed by a list of all ontology identifiers.

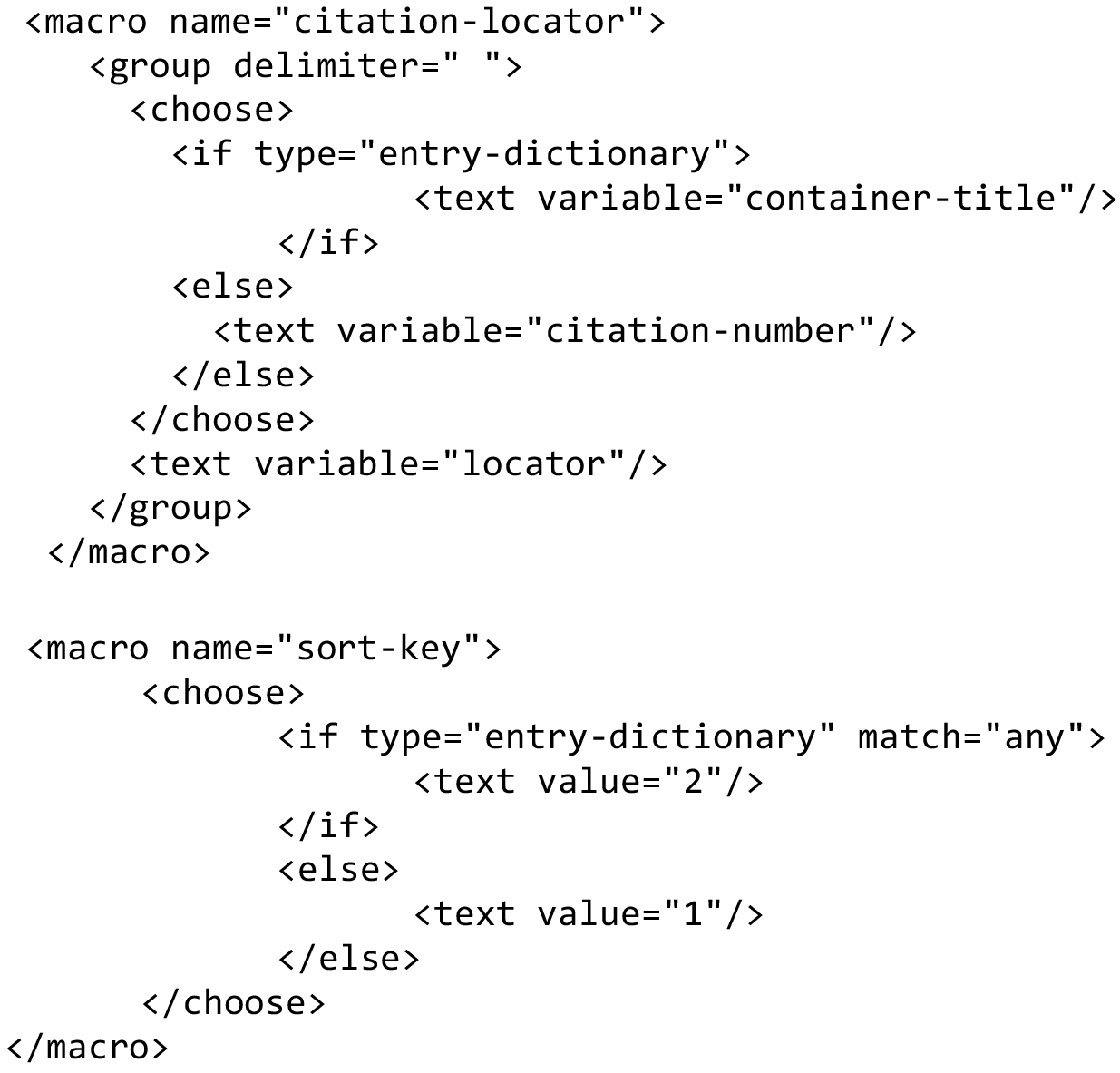

### Adapt citation block

The macro which correctly displays the ontology identifiers as citation labels in the text has to be called from within the citation block of the template csl-file. In the case of the Nature csl-file, we simply replaced the variable ‘citation-number’ and added a group-delimiter as exemplified here:

### Original file

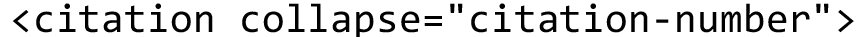

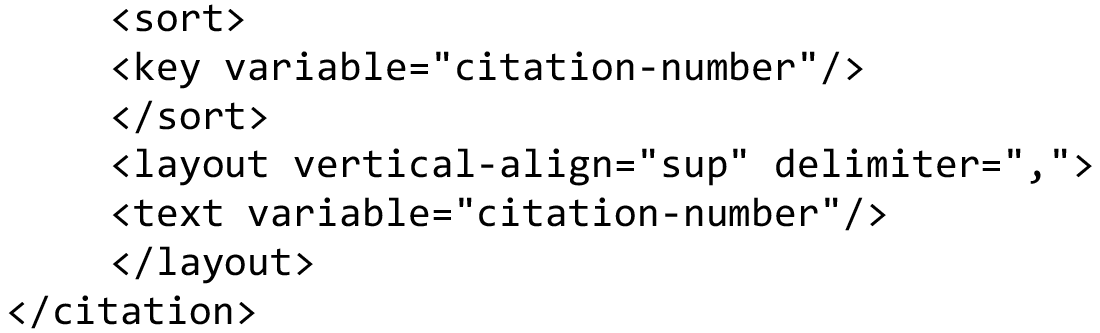

### Altered Phenotero-file

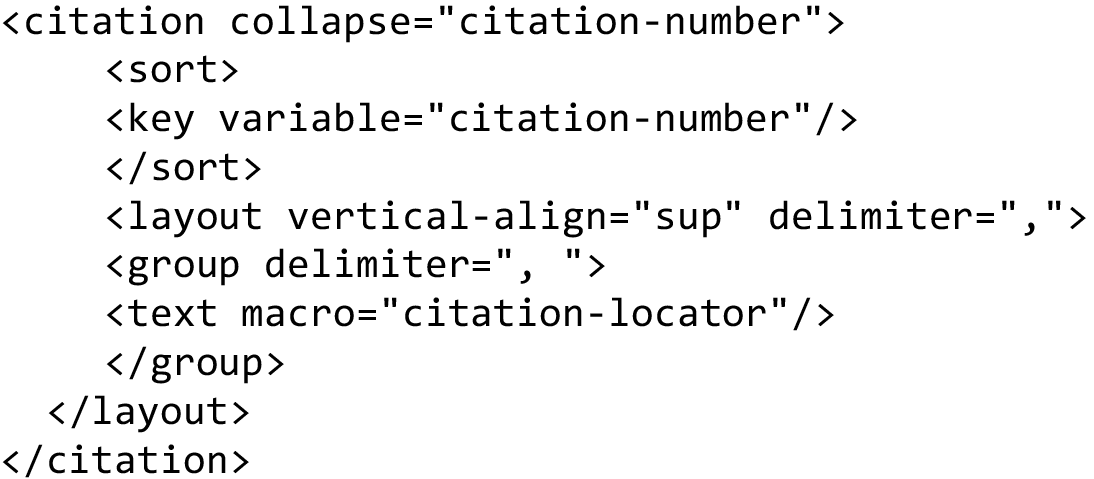

### Alter Bibliography section

The sorting macro has to be called from within the bibliography section to ensure that ontology identifiers and bibliography references are displayed separately. We added the call right at the start of the bibliography section as displayed here:

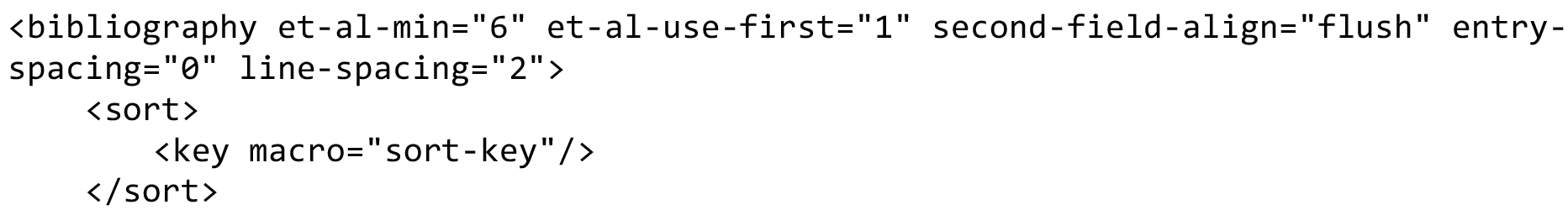

In addition, the program has to distinguish between *Phenotero* items and normal references. As *Phenotero* lists its items as ‘entry-dictionary’, we set up a separate condition for these items in the layout-part of the bibliography:

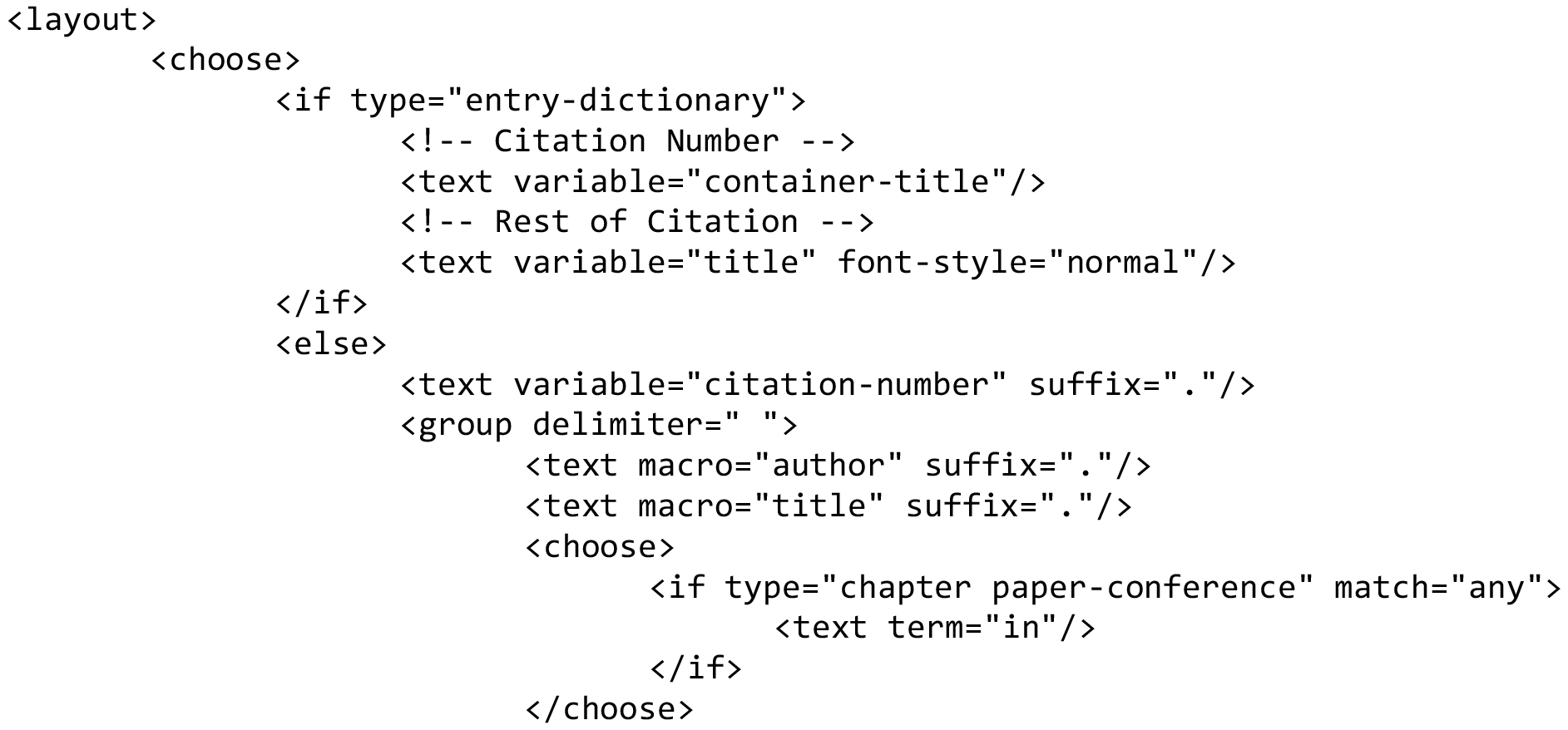

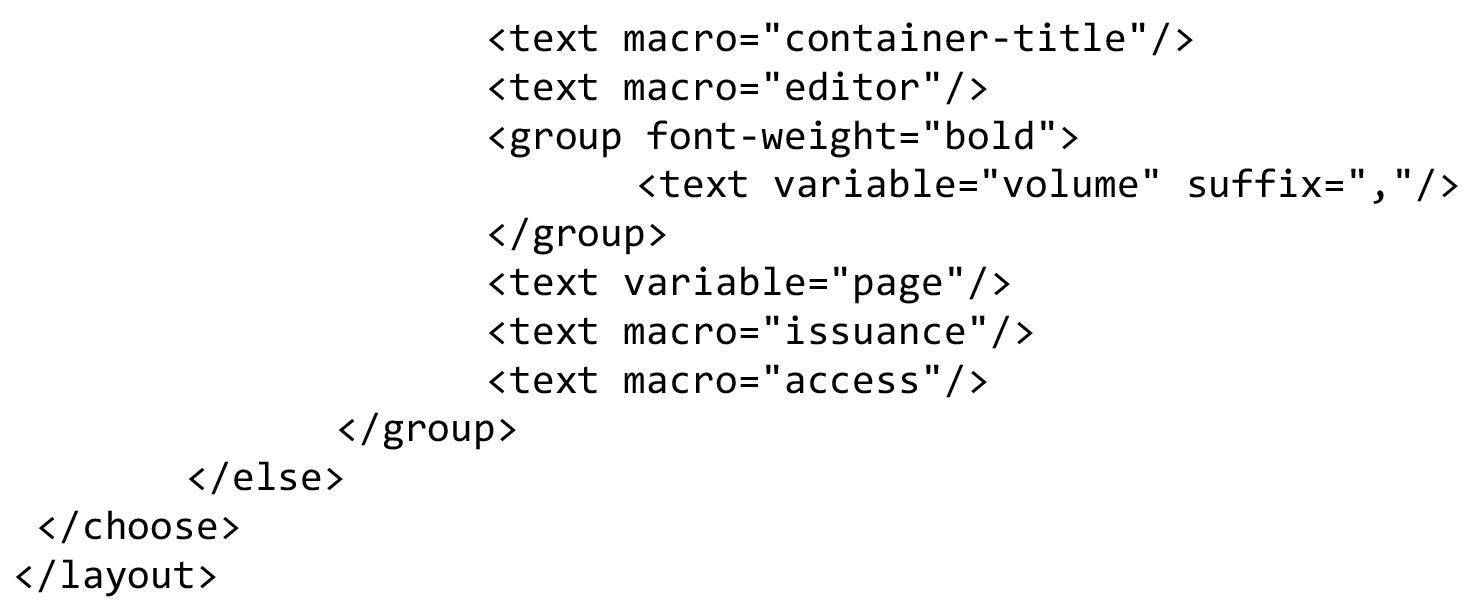

